# Pathway and network embedding methods for prioritizing psychiatric drugs

**DOI:** 10.1101/728055

**Authors:** Yash Pershad, Margaret Guo, Russ B. Altman

## Abstract

One in five Americans experience mental illness, and roughly 75% of psychiatric prescriptions do not successfully treat the patient’s condition. Extensive evidence implicates genetic factors and signaling disruption in the pathophysiology of these diseases. Changes in transcription often underlie this molecular pathway dysregulation; individual patient transcriptional data can improve the efficacy of diagnosis and treatment. Recent large-scale genomic studies have uncovered shared genetic modules across multiple psychiatric disorders—providing an opportunity for an integrated multi-disease approach for diagnosis. Moreover, network-based models informed by gene expression can represent pathological biological mechanisms and suggest new genes for diagnosis and treatment. Here, we use patient gene expression data from multiple studies to classify psychiatric diseases, integrate knowledge from expert-curated databases and publicly available experimental data to create augmented disease-specific gene sets, and use these to recommend disease-relevant drugs. From Gene Expression Omnibus, we extract expression data from 145 cases of schizophrenia, 82 cases of bipolar disorder, 190 cases of major depressive disorder, and 307 shared controls. We use pathway-based approaches to predict psychiatric disease diagnosis with a random forest model (78% accuracy) and derive important features to augment available drug and disease signatures. Using protein-protein-interaction networks and embedding-based methods, we build a pipeline to prioritize treatments for psychiatric diseases that achieves a 3.4-fold improvement over a background model. Thus, we demonstrate that gene-expression-derived pathway features can diagnose psychiatric diseases and that molecular insights derived from this classification task can inform treatment prioritization for psychiatric diseases.

## 1. Introduction

450 million people suffer from mental health disorders worldwide, with approximately 20% of Americans experiencing a mental illness in a given year.^1^ Additionally, mental illness negatively impacts quality of life, as it accounts for 32.4% of the years lived with disability.^2^ Even though most psychiatric drugs show benefits over placebo in clinical trials, nearly 75% of psychiatric prescriptions do not successfully treat the patient’s condition.^3,4^ Low drug efficacy for individual patients suggests a critical need to improve treatment prioritization for psychiatric disorders.

One reason for the low success rate is that gene-based biological mechanisms for disease are not routinely incorporated into clinical decision-making. Understanding the molecular basis of psychiatric diseases is necessary for rational drug choice. While traditional psychiatric diagnoses rely on feature sets composed largely of shared symptoms,^5^ recent research has identified some genetic contributions and has implicated the dysregulation of molecular pathways in the pathogenesis of these diseases.^6,7^ Gene expression levels provide quantitative measures to characterize these disrupted molecular pathways. For example, transcription studies have found that psychiatric diseases including schizophrenia (SCZ), bipolar disorder (BPD), and major depressive disorder (MDD) involve gene expression changes in neocortical regions responsible for cognitive and emotional control.^8^ The associated pathways may share some higher level characteristics, leading to similar symptoms, but probably also differ at a molecular level.^9^ Incorporation of the genetics of these complex diseases into clinical decision-making may allow tailored treatment options that more effectively modulate pathway-level disruption in the cell.

Recently gathered genomic datasets, ranging from single nucleotide polymorphisms (SNPs) for genome-wide association studies (GWAS) to gene expression data from RNA sequencing of brain tissue, provide a promising means for uncovering underlying causes of mental health disorders.^5^ Most recently, the PsychENCODE Consortium analyzed samples from over 2,000 individual brains and published several studies on gene expression and genomic regulation demonstrating shared genetic modules between multiple psychiatric disorders.^7,10,11^ These shared sets of genes make defining (and diagnosing) specific psychiatric diseases challenging. Therefore, effective diagnosis may require integration of cellular and molecular data to distinguish and classify psychiatric diseases.

Protein-protein interaction (PPI) networks, where nodes are proteins (i.e., gene products) and edges represent interactions between the proteins, are a useful framework for understanding disease at a molecular level.^12^ Previous work has analyzed gene expression data to find novel disease-gene associations and has even predicted non-psychiatric diseases via PPI networks and pathway-based approaches.^13^ The advantage of these network approaches is that they maintain a higher degree of biological interpretability compared to models using gene expression data directly as features. The PROPS^14^ algorithm was initially developed to classify types of inflammatory bowel disease and uses patient-specific pathway scores from gene expression data to evaluate pathways in the Kyoto Encyclopedia of Genes and Genomes (KEGG) resource. We hypothesize that network-based approaches such as PROPS may enable us to find distinguishing pathway features across psychiatric diseases.

Finding unique pathway features for these diseases may be useful for choosing drug treatments. At a molecular level, this task can consist of finding similar drug and disease gene signatures, on the assumption that drugs modulating disease-specific pathways may be more effective. In a network, a group of connected genes likely associated with the same disease are considered disease modules. Similarly, a group of connected genes likely known to be targets of a drug are drug modules. Proximity-based methods determining distance between drug and disease modules have been used to predict drug indications for a disease.^15^ However, embedding-based machine learning methods have recently achieved great success.^12^ For example, recent work has applied node2vec methods to create embeddings of PPI networks, thereby representing the network as feature vectors for each gene.^16,17^ Node2vec maximizes the likelihood of preserving neighborhoods of genes and structural features of the network. By doing so, it can capture complex relationships between nodes in lower dimensional space. Analyzing these feature vectors requires less computational power than performing expensive search and path-finding operations on the network itself. Thus, it has applications for multiple bioinformatics problems, such as predicting protein function, discovering novel protein-protein interactions, and finding new disease-associated genes.^18^

In this study, we use patient-specific gene expression data first to classify psychiatric diseases and find pathways that distinguish the diseases. Then, we use disease gene signatures and drug gene targets to recommend a prioritized list of disease-relevant drugs to patients (Fig. 1). We first create KEGG pathway scores from patient gene expression data and then used the pathways as features to build a classifier for psychiatric diseases. We define disease modules and drug modules by combining differential expression, literature-mining, and expert-curated databases. Using node2vec PPI network embeddings to assess proximity between drug and disease gene modules, we generate a ranking of drug indications for a disease of interest. Our work establishes the feasibility of using biological mechanism to drive both the diagnosis of psychiatric disease and the prioritization of treatments using patient-specific gene expression data.

**Figure 1:**
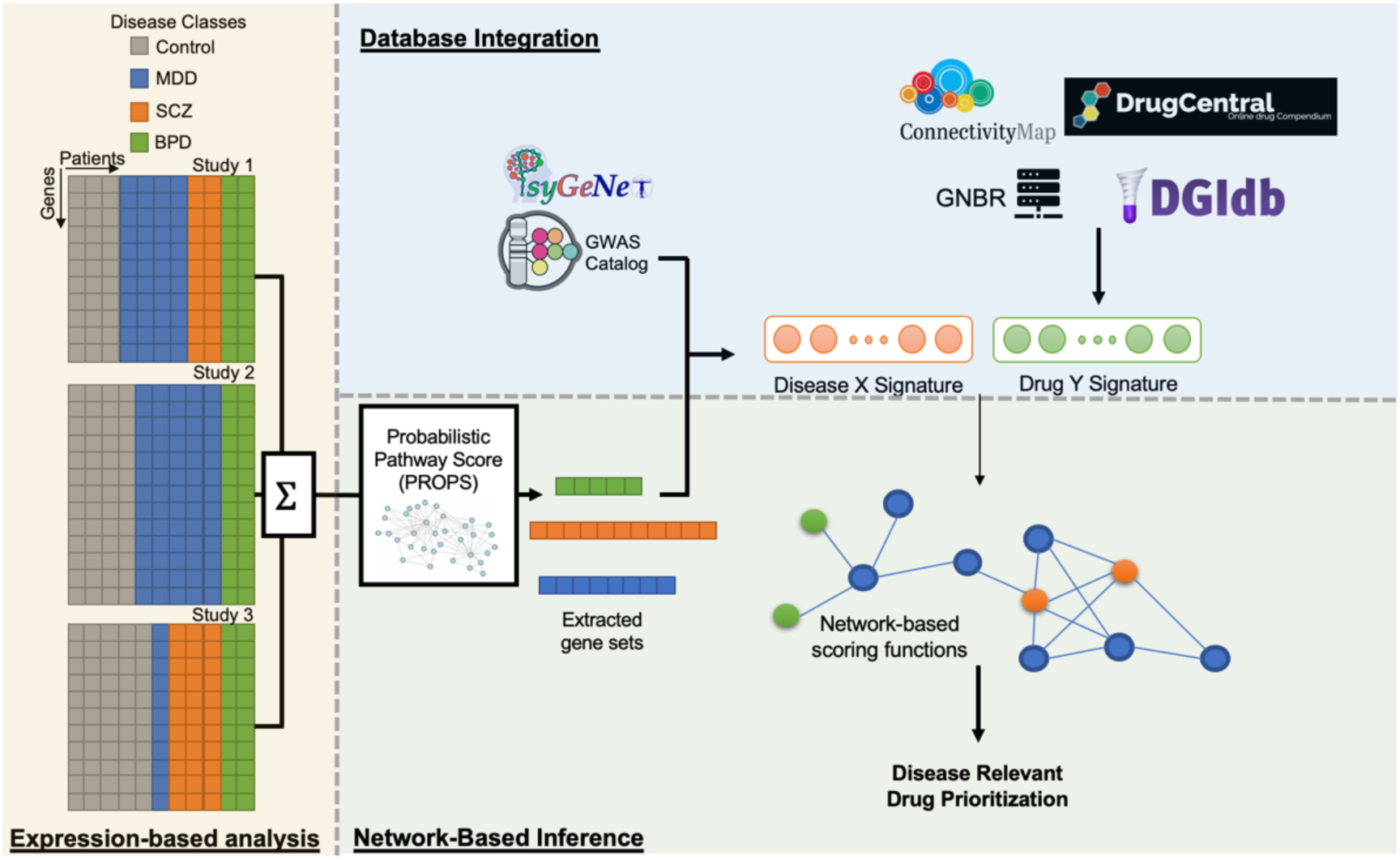
Integrated approach towards drug prioritization for psychiatric disease treatment. We partition our methods as expression-based analysis of transcriptome data from GEOdatabase integration of expert and experimentally-derived disease and drug gene signatures, and network-based inference techniques.

## 2. Methods

### 2.1. Collecting and processing gene expression data

In order to predict psychiatric diseases from gene expression data, we curated five datasets containing RNA expression data from SCZ, MDD, and/or BPD along with matched controls from the Gene Expression Omnibus (GEO): GSE92538^19^, GSE98793^20^, GSE27383^21^, GSE120340^22^, and GSE21138^23^. All five studies used Affymetrix microarrays to obtain expression data. We queried expression sets using the GEOquery package in R. Genes that mapped to multiple probes were averaged, as previously done.^14,24^ Next, using the genes common to all microarray platforms, we normalized the data using the affy package^25^ and corrected for batch effects using ComBat from the sva package.^26^ Overall, we processed expression data from 145 cases of SCZ, 82 cases of BPD, 190 cases of MDD, and 307 shared controls (Table 1).

**Table 1:** Summary of GEO studies queried for analysis. Information includes tissue of origin for each study (WB=whole blood, PFC=prefrontal cortex), along with number of cases and controls (SCZ=schizophrenia, BPD=bipolar disorder, MDD=major depressive disorder).

| Study | Tissue Type | # Control | # SCZ | # BPD | # MDD |
| --- | --- | --- | --- | --- | --- |
| GSE92538 | PFC | 175 | 62 | 62 | 62 |
| GSE98793 | WB | 64 | 0 | 0 | 128 |
| GSE27383 | WB | 29 | 43 | 0 | 0 |
| GSE120340 | PFC | 10 | 10 | 20 | 0 |
| GSE21138 | WB | 29 | 30 | 0 | 0 |
| Totals | PFC + WB | 307 | 145 | 82 | 190 |

### 2.2. Predicting disease from processed expression data

We evaluated the ability of unsupervised learning techniques, principal component analysis (PCA) and uniform manifold approximation and projection (UMAP), to distinguish samples by psychiatric disease using processed gene expression data. Unsupervised learning methods were evaluated based on visualization of feature vectors in the reduced 2-dimensional space.

Pathway importance scores for the individual samples were generated via the PROPS method.^14^ By using a random-walk-based approach, PROPS models KEGG pathways as Bayesian networks and returns a probabilistic pathway score that reflects pathway activity for each of the 268 KEGG biological pathways. We trained decision tree, random forest, and support vector classifiers from scikit-learn^27^ to distinguish psychiatric diseases from one another using pathway scores as features. Performance of the models was evaluated based on the micro average area under the receiver operating characteristic (auROC) across the three diseases using an iterative cross validation approach. We performed a grid search to tune hyperparameters of the classifiers. We determined the most important pathway features of the random forest model and inferred a gene importance score based on the frequency by which the genes associated with high scoring pathways.

### 2.3. Ranking disease-relevant drugs

After we performed disease prediction and identified the most important features (i.e., KEGG pathways), we sought to find disease-relevant drugs for psychiatric diseases from gene signatures associated with psychiatric diseases and gene targets associated with psychiatric drugs.

#### 2.3.1. Curating disease gene signatures

Initial disease signatures, defined as gene sets associated with psychiatric diseases, were procured from the Psychiatric disorders and Genes association NETwork (PsyGeNET), a database curated by domain experts which integrates genes from DisGeNET and literature-mining.^28^ The disease classes that we obtained from PsyGeNET were bipolar disorder, major depressive disorder, and schizophrenia, which are the three most common psychiatric disorders. Subsequently, we augmented these gene sets with genes linked to significant SNPs from GWAS Catalog for each respective disease.^29^

In order to validate these gene sets and potentially compare the genes associated with different psychiatric diseases, we performed functional analysis on the disease gene clusters. We characterized these gene sets for each disease by finding enrichment of the gene sets compared to all genes for specific molecular functions in the Gene Ontology (GO) database and Reactome pathway database with the clusterProfileR package.^30^ Reactome was chosen as an orthogonal pathway information source to validate findings based on KEGG-derived pathway scores. We visualized enrichment scores for each significant molecular function and disease to identify potential disease specific molecular mechanisms.

#### 2.3.2. Curating drug gene targets

We created a list of 275 pertinent psychiatric drugs through the Anatomical Therapeutic Chemical Classification (ATC) system, a hierarchical drug classification system which classifies drugs by tissue-specific therapeutic effects.^31^ To generate a list of drugs potentially relevant to BPD, MDD, and/or SCZ, we included all drugs classified as nervous system drugs (class N). The classes of these drugs included psycholeptics, psychoanaleptics, anti-Parkinson’s drugs, antiepileptics, analgesics, anesthetics, muscle relaxants, and other nervous system drugs.

For each potential psychiatric drug, we found gene targets based on expertly curated lists, literature-mining, and gene expression signatures. Databases from DrugCentral^32^ and the Drug-Gene Interaction Database (DGIdb)^33^ contain lists of target genes by drug. The global network of biomedical relationships (GNBR^34^), a knowledge graph derived from PubMed abstracts, contains chemical-gene relationships. Connectivity Map (CMap) from the Broad Institute^35^ includes expression data of multiple cell lines before and after treatment with small-molecule compounds. We used CMap to extract differentially expressed genes for each relevant perturbagen (abs(score) > 8) and added these genes to the sets of drug targets for each gene. Each drug has a median of 134 related genes with an interquartile range of 26.

#### 2.3.3. Recommending drugs for a disease

We used our curated drug and disease gene signatures to create a method that recommends disease-relevant drugs in a ranked manner. For each disease-specific recommendation, we operate under a common assumption^36^ that drugs indicated for a disease will modulate the disrupted pathways of the disease. We compared seven methods of evaluating disease-drug “similarity”:

(1) “random” method: While not technically a method, we report the proportion of drug indications present in our overall list indicated for a particular disease of interest.

(2) “simple overlap” method: We determined the fraction of genes intersecting and the Jaccard similarity between each drug and disease module. For each disease, we ranked drugs by the fraction of intersecting genes and Jaccard similarity. This method is the simplest method of comparison and does not account for any pathways or interactions between proteins. Subsequent methods used PPI networks to attempt to capture these interactions.

(3) “connected components” method: We calculated the shortest path between each pair of genes in the disease module in the STRING v10 PPI network (String)^37^ and included the genes along each path in a larger disease module. We then computed the Jaccard similarity between the expanded disease module and the drug target module.

(4) “mean path length” method: We found the shortest path length in String between every pair of genes with one from the disease module and one from drug module and then averaged each path length for a drug. A smaller path length indicates that the genes are closer in the PPI network and that it is more likely for changes in one gene to affect the other.

(5-7) network embeddings methods: Along with these simpler methods, we used node2vec embeddings to capture structural features of the PPI networks and to inform our drug ranking lists by disease. We performed node2vec embeddings on String and GNBR. (5) “String 32D” uses the node2vec embedding of String into 32-dimensions, and (6) “String64D” does into 64-dimensions. (7) “GNBR32D” uses the node2vec embedding of GNBR in 32-dimensions. After generating feature vectors for each gene node, we found the mean cosine similarity of each pairwise combination of genes from disease module and the drug module.

Each of these methods produces a ranked list of drugs for a given disease. We evaluate these methods by extracting the top 25 scoring drugs, roughly 10% of the possible drugs, and determined whether they were indicated for the disease of interest. Indications were derived from the Side Effect Resource (SIDER)^38^, a database built by from natural language processing of drug package inserts. We reported the percentage of drugs in the top 25 indicated for each disease.

#### 2.3.4. Augmenting disease modules with disease pathways important for diagnosis

We used patient-specific gene expression information to improve disease signatures for improving treatment prioritization. From the 20 most important KEGG pathways from our random forest models (as described in 2.2) for distinguishing BPD, MDD, or SCZ, we extracted genes belonging to these pathways and added them to our initial disease signatures. Genes were added to the disease signature if they were not used to diagnose other diseases. To test these augmented modules, we generated a prioritized list of drugs using this augmented module and compared the percentage of drugs indicated in the top 25 of the ranked list. To ensure that simply adding more genes does not increase the percent indicated, we added 250 random nodes from String to the disease sets and evaluated the top 25 drugs for these new randomly enlarged gene sets. We derived biological insight about our recommendations by measuring functional enrichment of Reactome pathways for the augmented disease modules, as described in 2.3.1.

## 3. Results

### 3.1. Predicting disease from processed expression data

To provide molecular insights into psychiatric disease, we created a method to predict disease from gene expression data. Because we combined data from multiple sources, we assessed tissue-source-specific or batch effects using sequential dimensionality reduction techniques, PCA and UMAP, to create two-dimensional vectors representing each sample.^39^ When we visualized these vectors, the samples did not form any clusters by batch or tissue-type, confirming that the batch correction was successful (Fig. 2A). However, when we used UMAP to visualize expression data by disease, we did not observe any clustering (Fig. 2B). This finding demonstrates the challenge of distinguishing psychiatric diseases from expression data.

**Figure 2:**
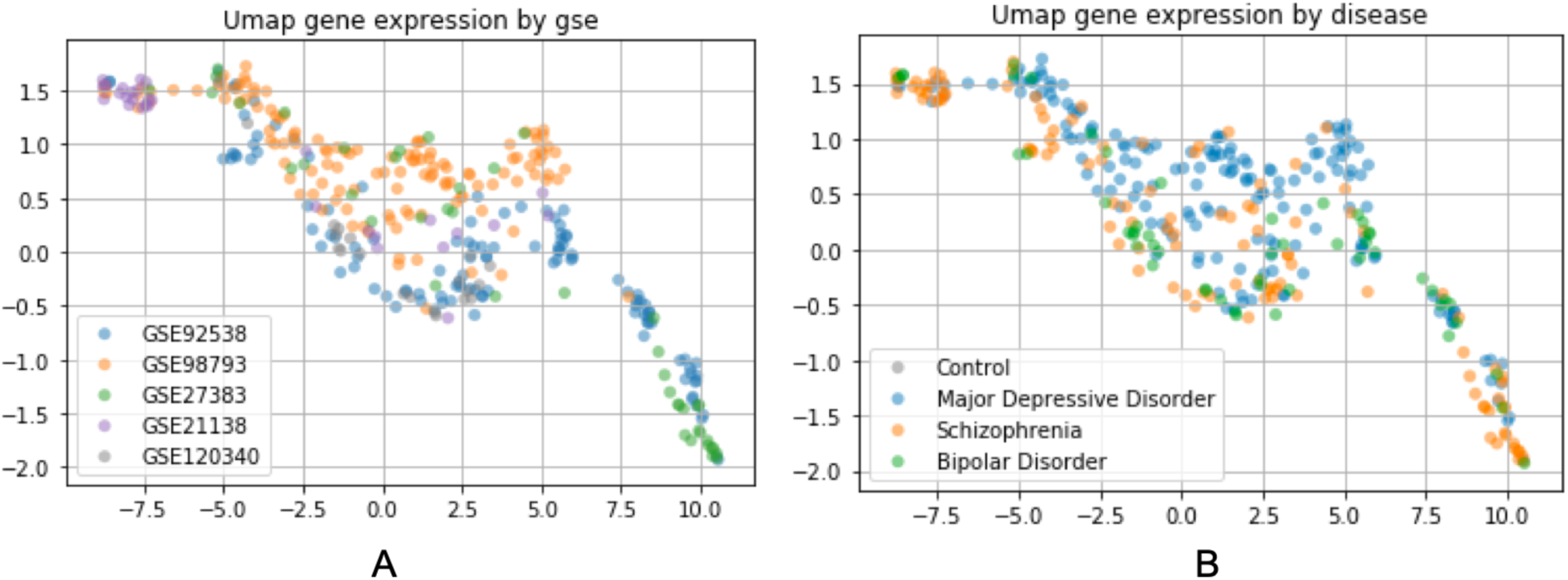
2-dimensional visualization after PCA and UMAP dimensionality reduction of batch-corrected gene expression data colored by (A) study and (B) psychiatric disease.

Pathway-informed classification approaches can overcome the noisiness in raw gene expression by aggregating signal as biologically meaningful features. We used the PROPS algorithm to generate new features (significant pathways), as opposed to raw expression values, for classifying the psychiatric diseases and providing insight into drug reprioritization. Fig. 3A shows ROC curves for the classification task for each disease, along with the micro and macro average ROC curves. We noted that the ROC curves for each disease are roughly similar indicating that our method is able to capture distinguishing features for all classes. After optimizing classifier parameters using via grid-search, we evaluated each multi-disease classifier using iterative cross-validation and report the auROC for the three methods (Fig. 3B). The random forest classifier performed the best, with an accuracy of 0.78 +/- 0.02. The support vector classifier performed second-best, followed by the decision tree classifier. For downstream feature selection, we used the random forest model because it performed the best. To minimize overfitting, we retrained on the classifier on the 50 most common features and observed no reduction in model performance. Therefore, using probabilistic KEGG pathway scores as features rather than raw expression data itself enables distinguishing of psychiatric diseases with accuracy.

**Figure 3:**
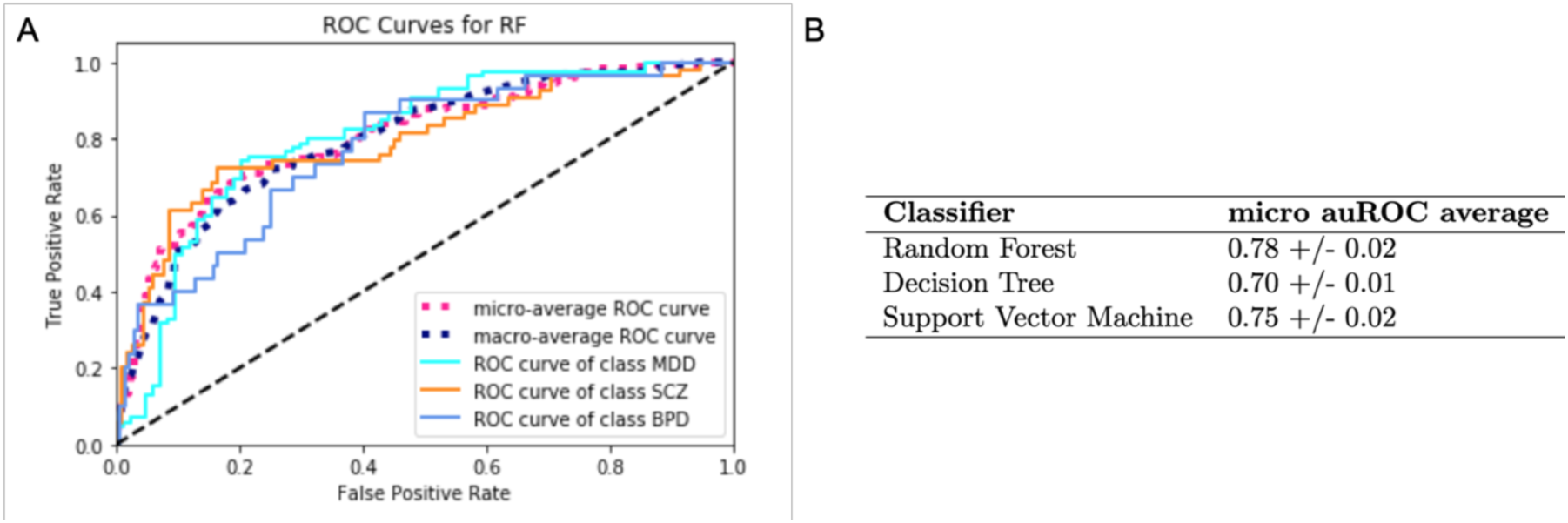
(A) ROC curve for a representative multi-class random forest (RF) model. (B) Micro auROC average +/- standard deviation shown for each classifier type across 100 cross-validation trials.

Additionally, probabilistic Bayesian methods can provide increased biological interpretability. For instance, KEGG pathways found to be important for schizophrenia according to our model include not only direct neurological disruption of GABA, cholinergic, and glutamatergic neurotransmitter signaling pathways, but also disruptions in folate metabolism. Folate deficiency has recently been shown in literature to be a risk factor for schizophrenia and its associated negative symptoms.^40^ Thus, pathway-based disease classification models can provide novel insights about the pathogenesis of these selected psychiatric diseases.

### 3.2. Ranking disease-relevant drugs

A synthesis of expert-curated sources (PsyGeNET, ATC, Drug Central, DGIdb), literature-mining databases (GNBR), and evidence from experiments (GWAS catalog, CMap) provided disease and drug modules for drug treatment prioritization. The integration of knowledge from multiple data sources may lead to valuable insight for therapeutic applications.

Our drug prioritization approach assumes that drugs should be more effective if their module has a greater degree of overlap with the corresponding disease module. Using the seven methods described in section 2.3.3, we show that the genetic overlap between drug and disease modules is significantly greater than random chance (Fig. 4A). We also note that embedding methods performed better than other methods such as “simple overlap” (21% were indicated in the top 25 drugs) or “mean path length” (27%). “String 32D” had an average accuracy across the three diseases of 48%, “String 64D” had an average of 45%, and “GNBR 32D” had an accuracy of 37%. The average indication for “random” was 15%. Embedding methods may better capture drug indications because they capture latent neighborhood structural features of the networks.

**Figure 4:**
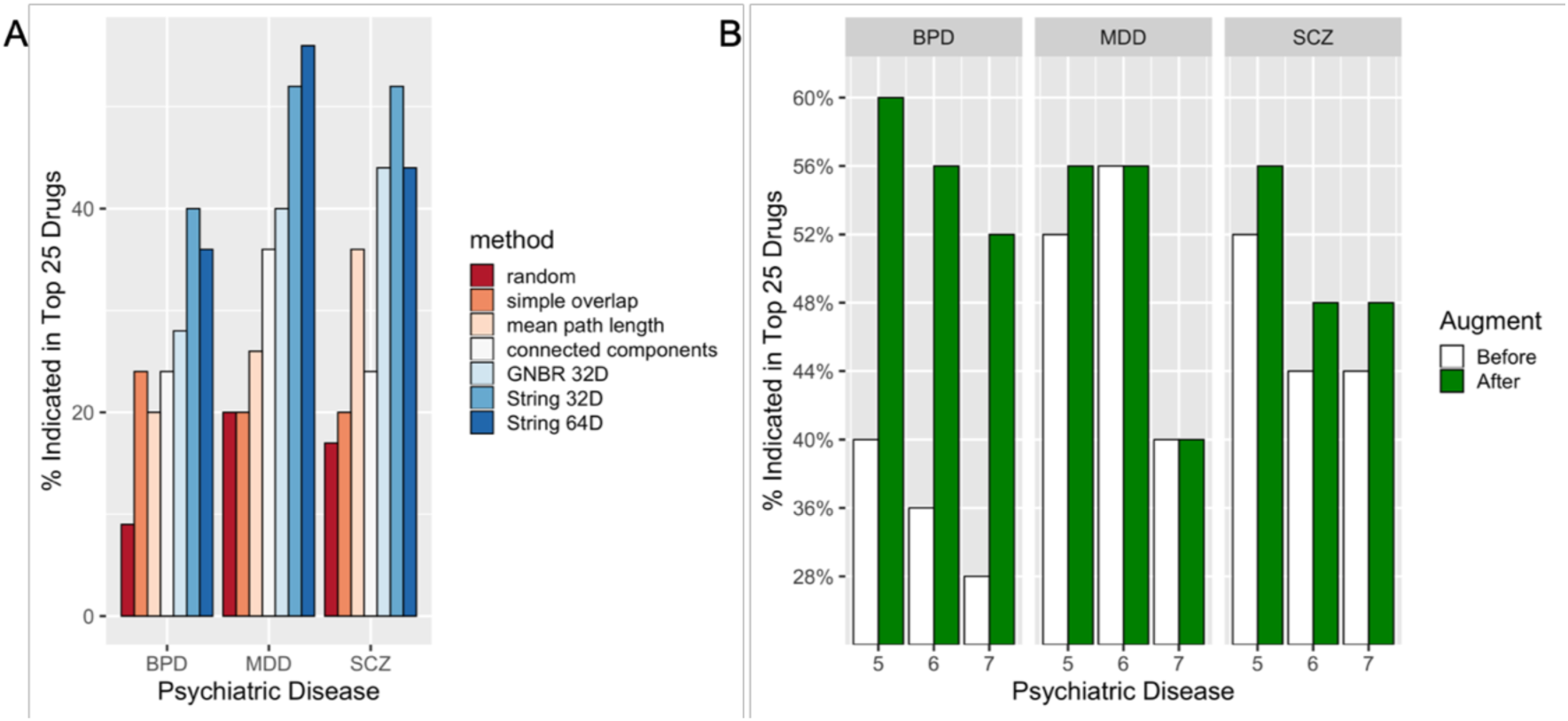
Evaluation of disease-relevant drug ranking methods clustered by disease and colored by method. (A) Original methods by disease. (B) Embedding methods with and without disease gene set augmentation. Numbers on the x-axis represent the embedding method used, 5 = String 32D, 6 = String 64D, and 7 = GNBR 32D.

One potential limitation of network embedding methods is that performance is dependent on the underlying network structure. Thus, they can be susceptible to incomplete data. For instance, most networks are unsigned and undirected, and thus the difference between a contraindication linkage and a treatment linkage is difficult to discern. As biological network data improves and our knowledge representation of biological mechanisms grows more complete, our embedding method will need to model these more complex genetic mechanisms.

We find that separating different psychiatric disorders is important for the prioritization of treatment options. Disease signatures augmented with pathway relevant genes perform disease diagnosis performed better than the initial disease signatures (Fig 4B). Given the accuracy of the pathway-based classification of psychiatric disease, this result is not necessarily surprising. However, it does emphasize the importance of pathway level dysregulation, as opposed to gene-level dysregulation in the treatment of psychiatric disorders.

We also note that adding additional genes to a curated gene set can affect performance in different ways. BPD had a large increase in capturing drug reprioritization targets, while the change for MDD was negligible. This is likely due to disease module for SCZ and BPD increased by approximately 200 genes each, while the MDD module changed by fewer than 20 genes. The increase in percentage indicated in the top 25 was not solely a result of adding any genes. When we added 250 random genes to the disease sets, the percentage indicated dropped to approximately 24% from over 50%. Even though there is a lack of novel genetic contributions, given that embedding methods for MDD performed well on the initial disease signature suggests that the expertly curated and literature derived gene sets may already be quite comprehensive.

While our drug indication labels represent only known drug-disease treatment pairs, we still can capture potentially novel targets. For example, niflumic acid is a pain reliever that was not present in the top indications list based on the original disease signatures but received a high score after the addition of 20 genes. While niflumic acid is not indicated for depression, it is believed, like other recently approved MDD treatments such as ketamine, to have antagonistic effects on the N-methyl-D-aspartate (NMDA) receptor.^41,42^ Thus, niflumic acid may serve as a promising MDD treatment candidate. Another repurposing example, chlorzoxazone, traditionally used as a muscle relaxant, was suggested as a drug relevant for schizophrenia. Recent research has implicated low activity of calcium-activated potassium (SK) channels as a potential pathway causing schizophrenia, and chlorzoxazone has demonstrated the ability to activate SK channels in mouse models.^43^ Thus, our method, with further experimental validation, may provide a means to suggest repurposable drugs with therapeutic effects.

Along with evaluating the performance of the new ranked list, we also used functional analysis to find enriched Reactome pathways and characterize the original and the augmented disease gene sets in order to elucidate differences between the psychiatric diseases. The original disease signatures showed extensive overlap in significantly enriched Reactome pathways, suggesting common mechanisms shared across the diseases (Fig. 5A). On the other hand, the enriched Reactome pathways for the augmented disease sets differed for some Reactome pathways (Fig. 5B). The differences can be attributed to the additional genes from the pathway-based diagnosis method, which emphasized distinct molecular pathways for the diseases. Namely, we observed that that Class C3 metabotropic glutamate/pheromone receptors distinguish SCZ from the other diseases and senescence-associated secretory phenotype (SASP) distinguished BPD from the others. Recent literature has implicated glutamate in SCZ pathogenesis in NMDA receptor hypofunction hypothesis, and some studies have discussed SASP as a marker of BPD.^44,45^ Thus, the Reactome enrichment demonstrated that the addition of modules from the pathway-based diagnosis method enabled easier disease delineation and drug prioritization.

**Figure 5:**
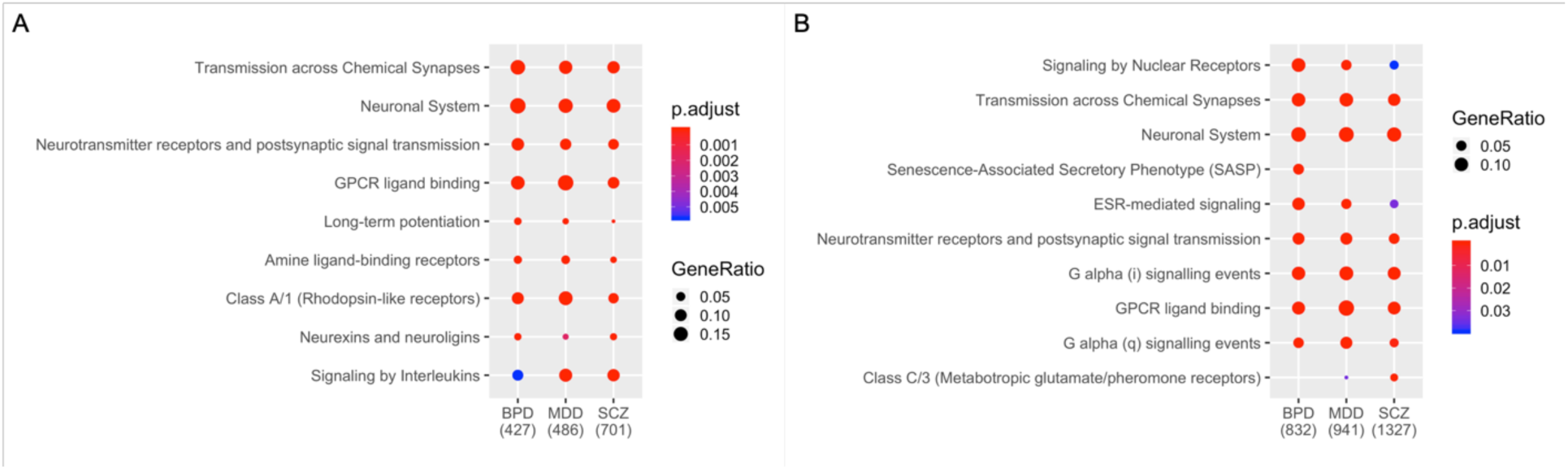
Dot plot of enriched Reactome pathways for (A) original gene sets for bipolar disorder (BPD), major depressive disorder (MDD), and schizophrenia (SCZ), (B) The same plot for augmented gene sets from disease-differentiating KEGG pathways (p > 0.05). We note the similarity of functional annotations in (A) and disease-specific differences in two pathways: SASP and Class C/3, in (B).

## 4. Conclusion

Psychiatric diseases are highly prevalent, and treatments are often unsuccessful. Lack of biological understanding of psychiatric disease pathogenesis, along with symptom-based, ambiguous diagnoses contribute to the low efficacy of psychiatric drugs. To address these challenges, we explored the use of gene expression data to classify the three main classes of psychiatric disorders: bipolar disorder, major depressive disorder, and schizophrenia. We subsequently used PPI networks and functional pathways to predict disease-relevant drugs and uncover molecular pathway distinctions between the three diseases. These enabled us to choose therapies that were more focused on disease-specific pathways.

In our work, we linked diagnostic separation with therapeutic prioritization. Specifically, we demonstrated that probabilistic pathway scores derived from gene expression data can find pathway differences between psychiatric diseases, enabling pathway-based analysis as a promising approach for both diagnostic and therapeutic exploration. We further refined the genetic linkages between drugs and psychiatric disease by curating drug and disease modules from a heterogeneous array of literature, expert, and experimental sources. Even with these curated gene sets, the informatic question of finding similarity between drug and disease modules is a complex task.

After comparing multiple methods, we found that node2vec embeddings of PPI networks best prioritized disease-relevant drugs from gene sets; these findings corroborate the idea that unsupervised embeddings capture latent structural features in networks. These structural features are not directly observable from the network itself. Additionally, when we augmented the baseline disease gene sets with genes from the most important KEGG pathways from our classifier for each disease, we observed an improvement in drug prioritization by disease. Functional analysis of these augmented disease gene signatures with orthogonal Reactome pathways demonstrated that these augmented disease signatures provide molecular pathway distinctions between the three psychiatric disorders studied here.

## Acknowledgements

Y.P is supported by the 2019 Undergraduate Research Summer Program in the Department of Bioengineering at Stanford University. M.G. is supported by the Medical Science Training Program Grant (5T32GM007365). RBA is supported by GM102365. Preprint of an article submitted for consideration in Pacific Symposium on Biocomputing.

## References

1. Steel, Z. et al. 1980-2013. Int. J. Epidemiol. 43, 476–93 (2014).

2. Vigo, D., et al. The Lancet Psychiatry 3, 171–178 (2016).

3. Seemüller, F., et al. BMC Med. 10, 17 (2012).

4. Amare, A. T., et al. EPMA J. 8, 211–227 (2017).

5. Goekoop, R. & Goekoop, J. G. PLoS One 9, e112734 (2014).

6. Tenenbaum, J. D. et al. Brief. Bioinform. 1–15 (2017).

7. Gandal, M. J. et al. Science (80-.). 362, eaat8127 (2018).

8. Gandal, M. J. et al. Science. 697, 693–697 (2016).

9. Geschwind, D. H. & Flint, J. Science 349, 1489–94 (2015).

10. PsychENCODE Consortium, T. P. et al. Nat. Neurosci. 18, 1707–12 (2015).

11. Li, M. et al. Science 80. 362, (2018).

12. Luo, P. et al. IEEE BIBM 1259–1264 (2016).

13. Ata, S. K. et al. BMC Syst. Biol. 12, 138 (2018).

14. Han, L. et al. Bioinformatics 34, 985–993 (2018).

15. Cheng, F., et al. Nat. Commun. 10, 1197 (2019).

16. Luo, P., et al. Bioinformatics (2019).

17. Grover, A. & Leskovec, J. Proceedings. Int. Conf. Knowl. Discov. Data Min. 855–864 (2016).

18. Zitnik, M., Agrawal, M. & Leskovec, J. (2016).

19. Hagenauer, M. H. et al. PLoS One 13, e0200003 (2018).

20. Leday, G. G. R. et al. Biol. Psychiatry 83, 70–80 (2018).

21. van Beveren, N. J. M. et al. PLoS One 7, e32618 (2012).

22. Abdolmaleky, H. M. et al. Am. J. Med. Genet. Part B Neuropsychiatr. Genet. (2019).

23. Narayan, S. et al. Brain Res. 1239, 235–248 (2008).

24. Davis, S. & Meltzer, P. S. Bioinformatics 23, 1846–1847 (2007).

25. Gautier, L., Cope, L., Bolstad, B. M. & Irizarry, R. A. Bioinformatics 20, 307–315 (2004).

26. Leek, J. et al. sva (2019).

27. Pedregosa, F. et al. J. Mach. Learn. Res. 12, 2825–2830 (2011).

28. Gutiérrez-Sacristán, A. et al. Bioinformatics 31, 3075–3077 (2015).

29. MacArthur, J. et al. Nucleic Acids Res. 45, D896–D901 (2017).

30. Yu, G., Wang, L.-G., Han, Y. & He, Q.-Y. Omi. A J. Integr. Biol. 16, 284–287 (2012).

31. WHO Collaborating Centre for Drug Statistics Methodology., (1993).

32. Ursu, O. et al. Nucleic Acids Res. 47, D963–D970 (2019).

33. Cotto, K. C. et al. Nucleic Acids Res. 46, D1068–D1073 (2018).

34. Percha, B. & Altman, R. B. PLoS Comput Biol 11, 1004216 (2015).

35. Lamb, J. et al. Science (80-.). 313, 1929–1935 (2006).

36. Ferrero, E. & Agarwal, P. BioData Min. 11, 7 (2018).

37. Szklarczyk, D. et al. Nucleic Acids Res. 47, D607–D613 (2019).

38. Kuhn, M., Letunic, I., Jensen, L. J. & Bork, P. Nucleic Acids Res. 44, D1075–9 (2016).

39. McInnes, L., Healy, J. & Melville, J. (2018).

40. El Mawella, et al. Egyptian Journal of Psychiatry 39.2 (2018): 89.

41. Dang, Yong-Hui, et al. (2014): 5151–5159.

42. Lerma, J. U. A. N., and R. Martin Del Rio. Molecular pharmacology 41.2 (1992): 217–222.

43. Imbrici, P., et al. Frontiers in genetics 4 (2013): 76.

44. Diniz, Breno Satler, et al. The American Journal of Geriatric Psychiatry 25.1 (2017): 64–72.

45. Maksymetz, J., et al. Molecular brain 10.1 (2017): 15.

